# Murine cytomegalovirus evolved a cell-cycle regulator (m54.5) within the highly conserved viral DNA polymerase gene

**DOI:** 10.1101/2025.08.04.668388

**Authors:** Yan Zheng, Vanda Juranic Lisnic, Stephanie Lamer, Andreas Schlosser, Lars Dölken, Manivel Lodha

## Abstract

Ribosome profiling (Ribo-seq) coupled with transcription start site profiling time-course analyses recently unveiled hundreds of novel viral gene products in lytic murine cytomegalovirus (MCMV) infection. One of these is the m54.5 open reading frame (ORF) located within the highly conserved viral DNA polymerase locus (M54). Interestingly, the m54.5 ORF is expressed from its own transcript (m54.5 RNA) with early gene expression kinetics, and at much higher levels than M54. In this study, we show that m54.5 encodes a nuclear viral protein (m54.5p) that contributes to cell cycle regulation during lytic MCMV infection. We show that m54.5p interacts with components of the anaphase-promoting complex/cyclosome (APC/C) and the phosphatase-6 (PP6) complex. Nocodazole mitotic arrest assays confirmed G1 cell cycle arrest and dysregulation by m54.5. Serum starvation revealed impaired cell cycle progression to S-phase. Notably, m54.5p is not conserved in other cytomegaloviruses but functionally mimics the UL21a protein of human cytomegalovirus (HCMV), which similarly targets the master cell cycle regulator APC/C to disrupt cell cycle progression. m54.5 thus represents convergent evolution to HCMV UL21a in MCMV within the highly conserved viral DNA polymerase gene. Nevertheless, we found that m54.5p is dispensable for viral replication in cultured mouse fibroblasts, indicative of redundant cell cycle regulation in lytic MCMV infection. These findings highlight a surprising genomic plasticity of herpesviruses, facilitating the evolution of an independent transcript encoding for a >200 aa gene product within a deeply conserved viral gene locus.

**Author Summary:** Systems biology approaches have revealed a surprising complexity of herpesvirus gene products. Using advanced sequencing approaches, we discovered a novel gene, *m54.5*, that independently evolved within a highly conserved region of the murine cytomegalovirus (MCMV) genome. This gene, which shows no conservation in other CMVs, produces a nuclear protein, m54.5p, abundantly expressed early during infection. We show that m54.5p interacts with host cell cycle regulators—the anaphase-promoting complex/cyclosome (APC/C) and phosphatase-6 (PP6)—to arrest cells in G1 phase and block progression into S phase. This function and underlying mechanism are reminiscent of the unrelated UL21a protein in human cytomegalovirus, illustrating how distinct viruses can evolve similar strategies to control host cell division. Despite its role in cell cycle disruption, m54.5p is not required for MCMV replication in cultured cells, suggesting redundant viral mechanisms. Our findings reveal an unexpected plasticity of herpesvirus genomes to evolve new, functional transcripts and proteins even within one of the most highly conserved genomic regions. Our findings thereby reshape our understanding of herpesvirus evolution and virus-host interaction.

## Introduction

Human cytomegalovirus (HCMV) is a double-stranded DNA virus with a genome of ∼240 kb, defining the beta-herpesvirus subfamily [1,2]. It establishes lifelong persistent infections in humans [3]. Globally, HCMV infection affects approximately 60% to 90% of the population, with higher prevalence in developing countries [4]. While often asymptomatic in healthy individuals, HCMV poses significant risks to immunocompromised patients, transplant recipients, and neonates [5]. In neonates, congenital HCMV infection is the leading cause of intellectual disabilities and sensorineural hearing loss [6]. The virus exhibits strict species specificity, which presents challenges in studying its biology and immune evasion mechanisms [7].

Murine cytomegalovirus (MCMV) possesses a genome approximately 230 kb in size, exhibiting significant similarity to the human cytomegalovirus (HCMV) genome, thereby serving as a valuable model for studying HCMV pathogenesis, immune evasion and control [8]. Both viruses have developed diverse mechanisms to invade host cells and establish infection, heavily relying on the host cellular machinery. Their genome replication is closely linked to the host cell cycle, which comprises the mitotic (M), growth 1 (G1), DNA synthesis (S), and growth 2 (G2) phases in actively dividing cells [9–12]. Although both viruses encode their own DNA polymerases (UL54 in HCMV; M54 in MCMV) and replication factors for viral DNA synthesis, they must also manipulate the host replication environment to secure resources such as deoxynucleotides (dNTPs) and replication factors [13,14]. This task is particularly challenging because most host cells infected in vivo are in a non-dividing, quiescent state (G0), offering a limited pool of nucleotides for viral genome replication. To overcome this, cytomegaloviruses evolved strategies to induce a pro-proliferative state in host cells, prompting re-entry into the cell cycle from G0. Subsequently, they establish an atypical S-like phase - also referred to as pseudo-S-phase – which promotes viral genome replication while simultaneously inhibiting cellular DNA synthesis by arresting the cell cycle at the G1/S transition [11,15–17].

Over the years, numerous mechanisms employed by both murine cytomegalovirus (MCMV) and human cytomegalovirus (HCMV) to induce G1/S phase arrest have been extensively documented [10–12,18,19]. Both viruses utilize similar pathways, including modulation of the cell cycle regulatory Rb-E2F transcription factors, p53-p21 checkpoint, the DREAM complex, as well as targeting cyclins A and E kinases and SAMHD1 proteins [20–28]. However, substantial differences between these two viruses exist at the mechanistic level, particularly in the viral proteins involved. This is exemplified by the MCMV kinase M97 and its HCMV homolog, UL97, which modulate the DREAM complex by targeting the LIN54 and LIN52 subunits, respectively [27,28]. MCMV M117 targets the E2F 1–5 transcription factors, whereas its HCMV counterpart, UL117, specifically targets the mini-chromosome maintenance (MCM) complex—most notably MCM2 and MCM4—blocking their accumulation and chromatin loading to inhibit host DNA replication [18,29]. The HCMV UL82 gene product, pp71, drives quiescent cells into the cell cycle through its LxCxE motif and promotes the proteasome-dependent degradation of hypophosphorylated retinoblastoma (Rb) protein and its family members, p107 and p130—an interaction not conserved in MCMV homologs, M82 and M83[30]. HCMV pp150 (pUL32) contains an RXL motif that binds cyclin A, restricting IE gene expression in S/G2 phases and preventing mitotic cell death in cooperation with pUL21a-mediated cyclin A degradation [25,31]. In contrast, MCMV lacks this regulation in its pp150 homolog but utilizes M97, which binds cyclin A to alter its function and localization [27]. In summary, cytomegaloviruses have evolved multiple viral proteins to manipulate the cell cycle to promote productive infection.

APC/C is an evolutionarily conserved E3 ubiquitin ligase that controls cell cycle progression by mediating the spatiotemporal ubiquitination and degradation of key regulatory proteins [2,32]. The pUL21a, which is conserved only in primate CMVs, binds directly to APC/C, specifically targeting its subunits APC4 and APC5 for degradation [32]. Degradation triggers the destabilization and dissociation of the APC/C, disrupting its E3 ubiquitin ligase activity and ultimately promoting cell cycle dysregulation to facilitate viral replication. Regulation of the APC/C complex by MCMV had not been reported [12].

Recently, we provided a comprehensive re-annotation of the MCMV genome, studying lytic MCMV infection in NIH-3T3 murine fibroblasts. Using ribosome profiling (Ribo-seq) and transcription start site profiling (cRNA-seq and dRNA-seq), we annotated 380 viral transcripts and 454 viral open reading frames (ORFs). The majority of so far unknown viral ORFs included short, upstream and upstream-overlapping ORFs (uORFs and uoORFs), consistent with recent findings for several herpesviruses, including HCMV [2]. However, we also identified 68 so far unknown large viral ORFs of >100 amino acids (aa) in size. One of these is m54.5 ORF, which is expressed from its own viral transcript (m54.5 mRNA), initiating within the M54 gene. The m54.5 ORF is thus located fully within the highly conserved viral DNA polymerase (M54) ORF but utilizes the M54 poly(A) signal (**Fig 1**). m54.5 was excluded in previous in-silico predictions due to its localization within the M54 ORF. In this study, we characterized the m54.5 ORF product (m54.5p). We show that m54.5p is an early nuclear protein important in inducing cell cycle arrest at the G1/S phase. m54.5p interacts with key cell cycle regulators, including the APC/C complex, to manipulate the cell cycle, and thus evolved as a functional ortholog of the HCMV pUL21a gene within the deeply conserved viral DNA polymerase gene.

**Fig 1:**
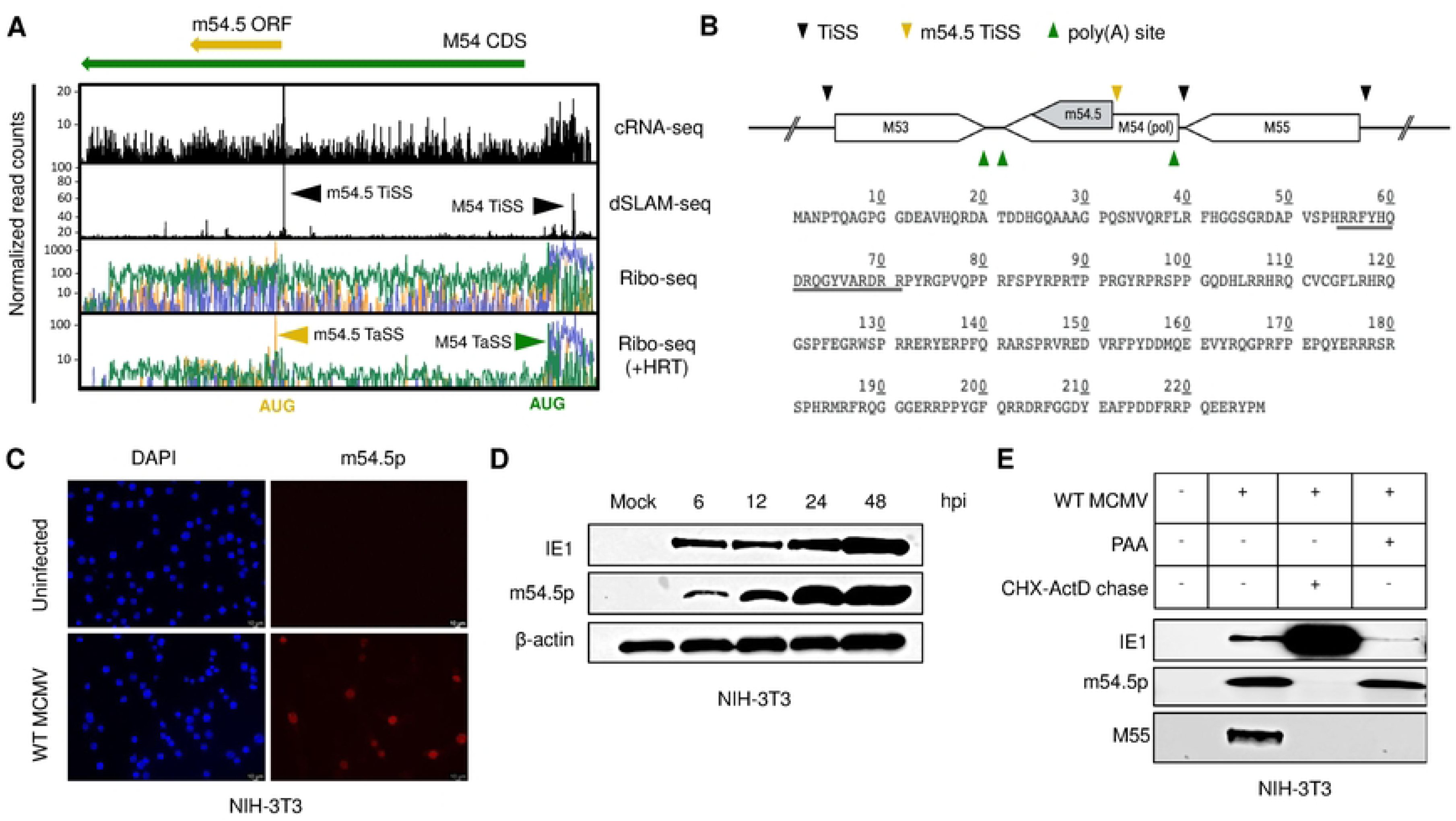
m54.5 encodes a nuclear m54.5 (m54.5p) expressed with early kinetics. **A**. Schematic representation of annotated ORFs and transcripts in the MCMV M54 and m54.5 loci. Normalized read counts (y-axis) for ribosome profiling (Ribo-seq), as well as transcription start site profiling data (dSLAM-seq and cRNA-seq) (8) aggregated across a time course during MCMV lytic infection in NIH-3T3 fibroblasts are shown. dSLAM-seq transcription start sites (TiSS) are depicted in black. Translation start sites (TaSS) are depicted in green and yellow for the M54 and m54.5 ORFs, respectively. Ribosome profiling data (including translation start site enrichment using harringtonine – HRT) are represented in logarithmic scale. The two transcription start site (TiSS) profiling approaches, cRNA-seq and dSLAM-seq, are represented on a linear scale. This schematic figure is based on data published by Lodha et al (8). **B**. Schematic diagram and amino acid sequence of the m54.5 ORF (227 aa). The bipartite NLS is underlined. The TiSS and poly-A sites of M53, M54, m54.5, and M55 are indicated. **C**. Uninfected NIH-3T3 cells and NIH-3T3 cells infected with WT MCMV at a multiplicity of infection (MOI) of 1 for 24 hours were subjected to immunostaining for m54.5p. DAPI was used for nuclear counterstaining. Data from two biological replicates are shown. **D**. Time-course immunoblot profiling of m54.5p during lytic MCMV infection of NIH-3T3 fibroblasts. Mock represents uninfected cells. Immunoblotting was additionally performed for MCMV IE1 and β-actin (loading control). Data from two biological replicates are shown. **E**. Kinetic analysis of m54.5p in MCMV-infected NIH-3T3 cells. Immunoblotting for various conditions shown was performed for m54.5p, IE1, and M55. Data from two biological replicates are shown. PAA: Phosphonoacetic acid, CHX: Cycloheximide, ActD: Actinomycin D.

## Results

### m54.5 encodes a nuclear protein (m54.5p) expressed with early kinetics

We previously conducted a comprehensive reannotation of the MCMV gene products expressed during lytic infection of murine NIH-3T3 fibroblasts [8]. This resulted in the identification of a novel ORF (m54.5), internal to the MCMV polymerase (M54 CDS), expressed at even higher levels than the viral DNA polymerase itself **(Fig 1A)**. BLAST analysis indicated that the m54.5 ORF is 100% conserved in all known MCMV strains without a single nucleotide mutation (**S1 File**). The absence of any significant stretch of codons without a stop codon in any other frame of M54 homologues in other cytomegaloviruses excludes conservation of m54.5 in other cytomegaloviruses and thus specifically evolved in MCMV (**S2 File**). Moreover, m54.5 lacks amino acid homology in other CMV species (including rat and human CMV) as well as UNIPROT cellular proteins (**S3 File**). The m54.5 gene thus independently evolved within MCMV. Subcellular localization prediction using PSORT II indicated a strong classical bipartite nuclear localization signal (NLS) **(Fig 1B)**, which contained an ^178^RSRSP^182^ sequence, previously shown to regulate nuclear localization of the RBM20 splicing factor [33]. Given its high expression levels and interesting location within the essential viral polymerase (M54), we decided to study the function of m54.5p.

Since m54.5 ORF could not be tagged in MCMV without affecting the viral polymerase, we generated a mouse monoclonal antibody against m54.5p. Immunofluorescence analysis of MCMV-infected cells at 24 hours post-infection (hpi) and in cells stably transduced for ectopic expression of m54.5p validated antibody specificity and confirmed the nuclear localization of m54.5p **(Fig 1C)**. Next, we performed a time-course analysis of m54.5p in MCMV lytic infection via immunoblotting. This revealed m54.5p to become readily detectable by 6 hpi, consistent with our previous data [8] **(Fig 1D)**. Chemical inhibition of viral DNA synthesis using phosphonoacetic acid (PAA) had no effect on m54.5p expression. To further characterize m54.5p expression kinetics, we performed a Cycloheximide (CHX)–Actinomycin D (ActD) reversal experiment. In this approach, CHX is first applied to block *de novo* viral protein synthesis while permitting transcription of immediate-early (IE) genes. Subsequent CHX removal and ActD addition allowed translation of newly transcribed viral IE mRNAs. These experiments confirmed that m54.5p is expressed with early kinetics (**Fig 1E**). In agreement with previous findings, these results validate m54.5p as a previously uncharacterized, 227-amino-acid nuclear viral protein encoded by the m54.5 ORF and expressed with early gene kinetics.

### An m54.5p null virus exhibits comparable growth and replication to wild-type MCMV

To elucidate the importance of m54.5p, we constructed an MCMV mutant (*m54.5mut* MCMV) on the pSM3fr backbone (*en passant* mutagenesis[34]) by mutating the start codon of m54.5 (AUG>UUG) without altering the M54 amino acid sequence **(Fig 2A)**. The newly generated mutant showed no m54.5p expression by Western blot and IF (**Figs 2B and 2C**). It is important to note that there are no downstream in-frame AUGs within 468 nt that would enable translation initiation and expression of a truncated m54.5p. Multistep growth curve analysis on NIH-3T3 fibroblasts revealed that the *m54.5mut* MCMV replicated with kinetics comparable to those of wild-type (WT) MCMV **(Fig 2D)**. We conclude that m54.5p is dispensable for viral replication and growth in cultured mouse fibroblasts.

**Fig 2:**
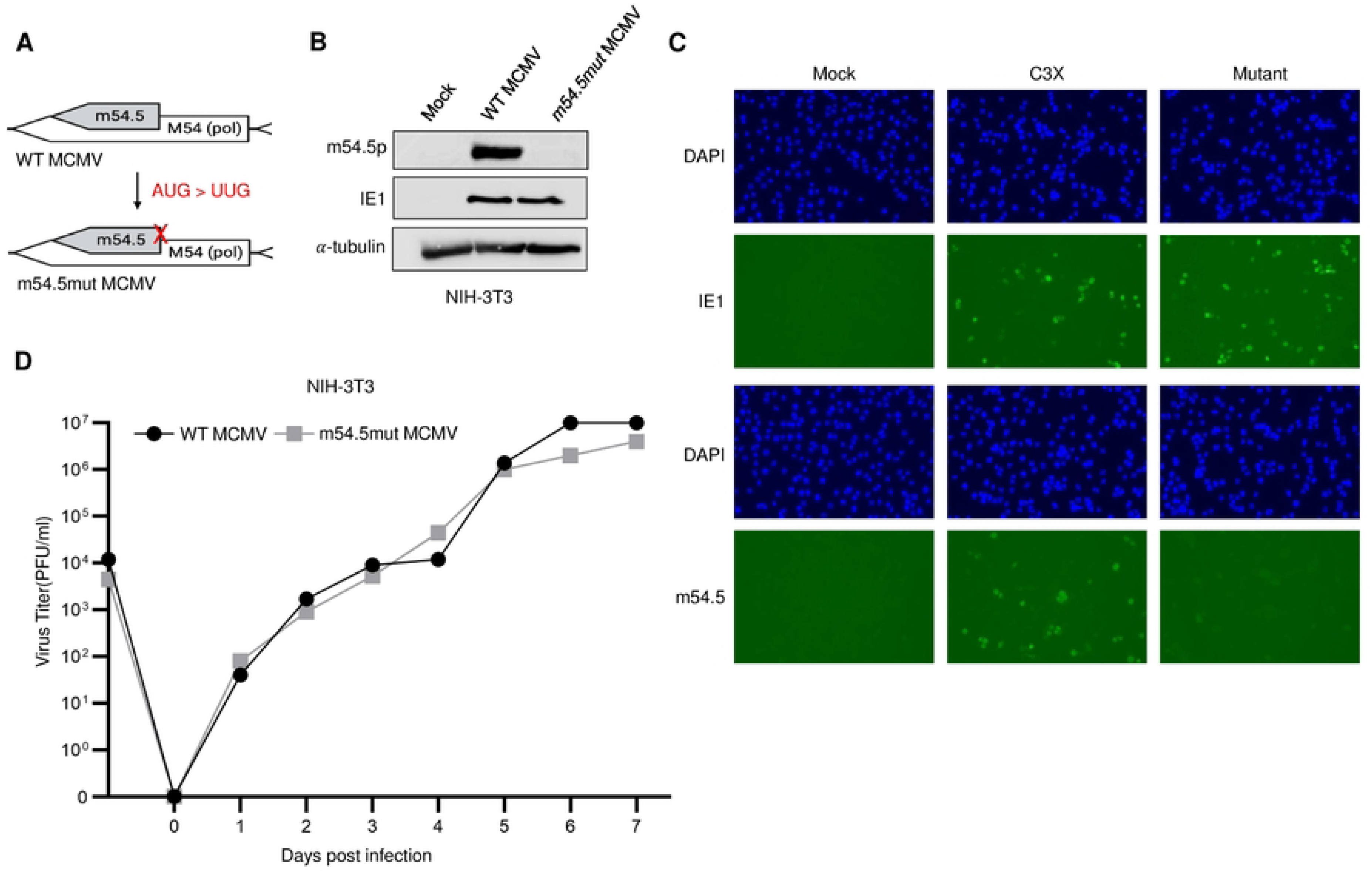
An m54.5p-null virus exhibits comparable growth kinetics to wild-type MCMV. **A**. Schematic representation of the mutagenesis approach for *m54.5mut* MCMV. **B**. NIH-3T3 cells infected for 24h with the indicated viruses at an MOI of 1 were immunoblotted for m54.5p, MCMV IE1, and α-tubulin (loading control). Mock represents uninfected NIH-3T3 cells. Data from two biological replicates are shown. **C**. NIH-3T3 cells infected with the indicated viruses at an MOI of 1 for 24 h were immunoblotted for DAPI, m54.5p, and IE1. **D**. Multistep growth curve analysis in NIH-3T3 cells. NIH-3T3 cells were infected with either WT MCMV or *m54.5mut* MCMV at an MOI of 0.1. Supernatants from infected cells were collected at the indicated time points post-infection (every 24 h) and titrated to determine viral yields. The experiment was performed in triplicate. The sample labeled ‘0.1’ represents the supernatant collected immediately after viral adsorption, followed by washing with 1x PBS.

### m54.5p interacts with components of the PP6 and APC/C complex

As m54.5p lacks homology to any known cellular or viral proteins, we resorted to functionally characterizing m54.5p by studying its interactome using co-immunoprecipitation (Co-IPs) combined with LC-MS. To this end, we generated polyclonal cell lines constitutively expressing C-terminally tagged m54.5p, m54.5p-V5-3T3, and m54.5p-FLAG-3T3 **(Fig 3A)**, where the latter cell line was utilized as a negative control for performing V5 tag Co-IPs. This ensured that any qualitative or quantitative changes in the cellular proteome due to m54.5p were accounted for. After validating the cell lines and analyzing V5 Co-IP efficiency **(Fig 3B)**, we performed Co-IP LC-MS for both cell lines under both uninfected and MCMV-infected (24 hpi) conditions (three biological replicates per condition). **Fig. 3C** shows the interactome of m54.5p in MCMV-infected samples plotted as a volcano graph depicting the enrichment (log_2_ fold change (V5/FLAG)) on the x-axis vs significance (adj. p-values) of the individual interactions on the y-axis. Top interactions were filtered according to p-values, and further assessment of interactors was performed by filtering for >4-fold enrichment. We then analyzed known protein complexes by filtering for high-confidence interactions via STRING (minimum score: 0.7/1). These revealed components of three distinct complexes, including the small subunit of the mitochondrial ribosomal proteins (MRPS), the protein phosphatase-6 complex (PP6), and the anaphase-promoting cyclosome complex (APC/C or ANAPC) **(Table 1)**. As m54.5p is a nuclear protein, we focused on proteins well-known to localize to the nucleus, which included the PP6 and APC/C complex. To validate our data, we confirmed interactions of m54.5p with the top interacting components of the respective complexes, including APC1 (ANAPC1), PP6, and PP6R3 by V5 Co-IP and immunoblotting **(Fig 3D)**.

**Fig 3:**
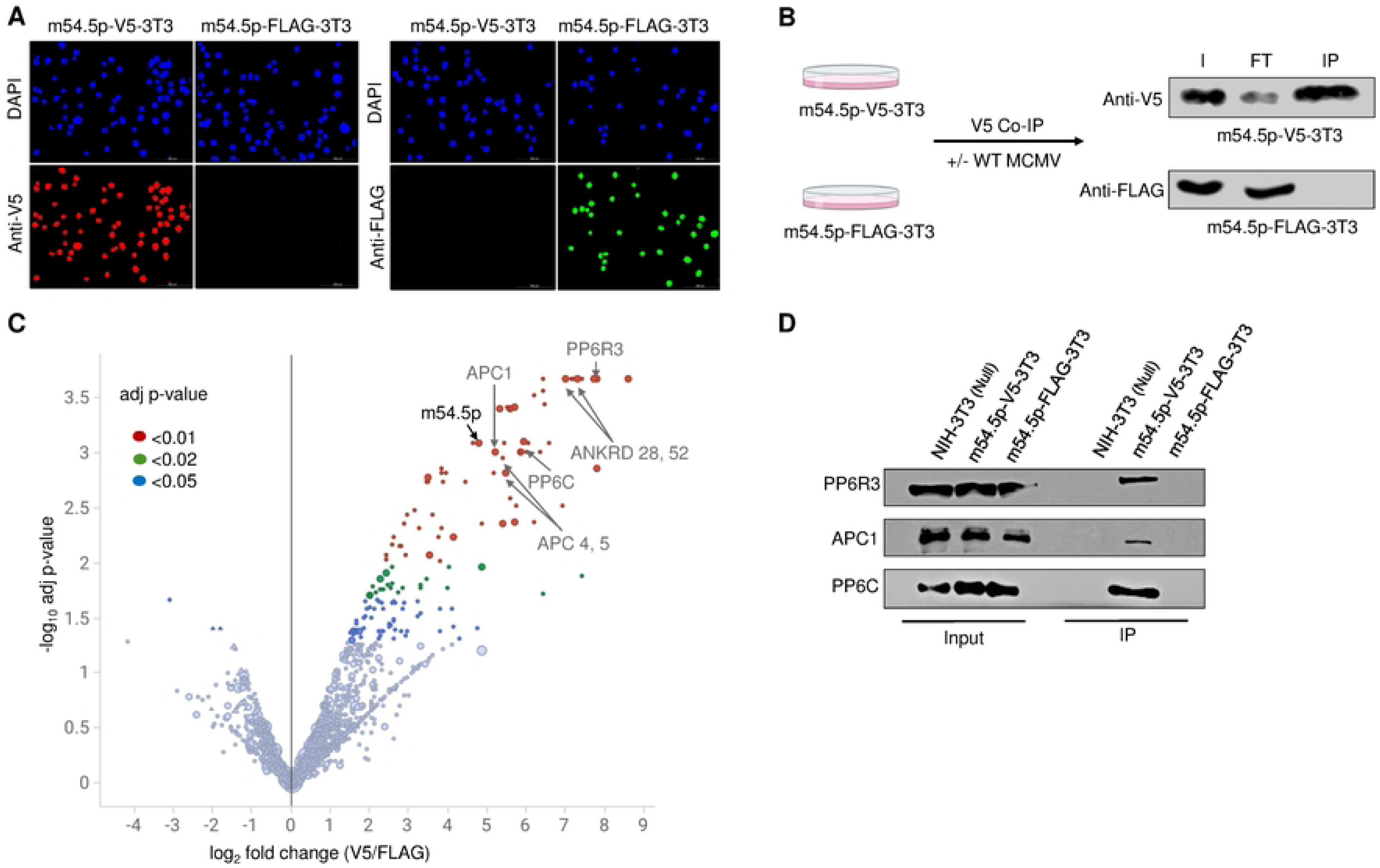
m54.5p interacts with components of the PP6 and APC/C complex. **A.** m54.5p-V5-3T3 and m54.5p-FLAG-3T3 cell lines were stained for V5 and FLAG tag epitopes. DAPI was used as nuclear counterstain. **B.** MCMV-infected (MOI=5, 24h) or uninfected m54.5p-V5-3T3 and m54.5p-FLAG-3T3 cells were subject to V5 Co-IP. Representative Western blots for the V5 Co-IP are shown. Efficiency is shown by enrichment of V5-tagged m54.5p in the IP sample. I: Input m54.5, FT: Flow-through after IP; IP: Co-IP Output. **C.** Volcano plot showing m54.5p interactome analysis by Co-IP LC-MS conducted for V5 Co-IP. The y-axis represents −log10 adjusted p-values for each interaction, while the x-axis depicts the log2 fold change representing the enrichment of V5 over FLAG. Colors represent varying degrees of statistical significance. Components of the PP6 (PP6R3, PP6C, ANKRD 28 and 52) and APC/C complex (APC 1, 4, and 5) are shown. Data from three biological replicates of MCMV-infected samples are shown. **D.** Immunoblot validation of m54.5p interactors (PP6R3, PP6C, APC1) by V5 Co-IP as performed in B. Data from two biological replicates are shown.

**Table 1:**
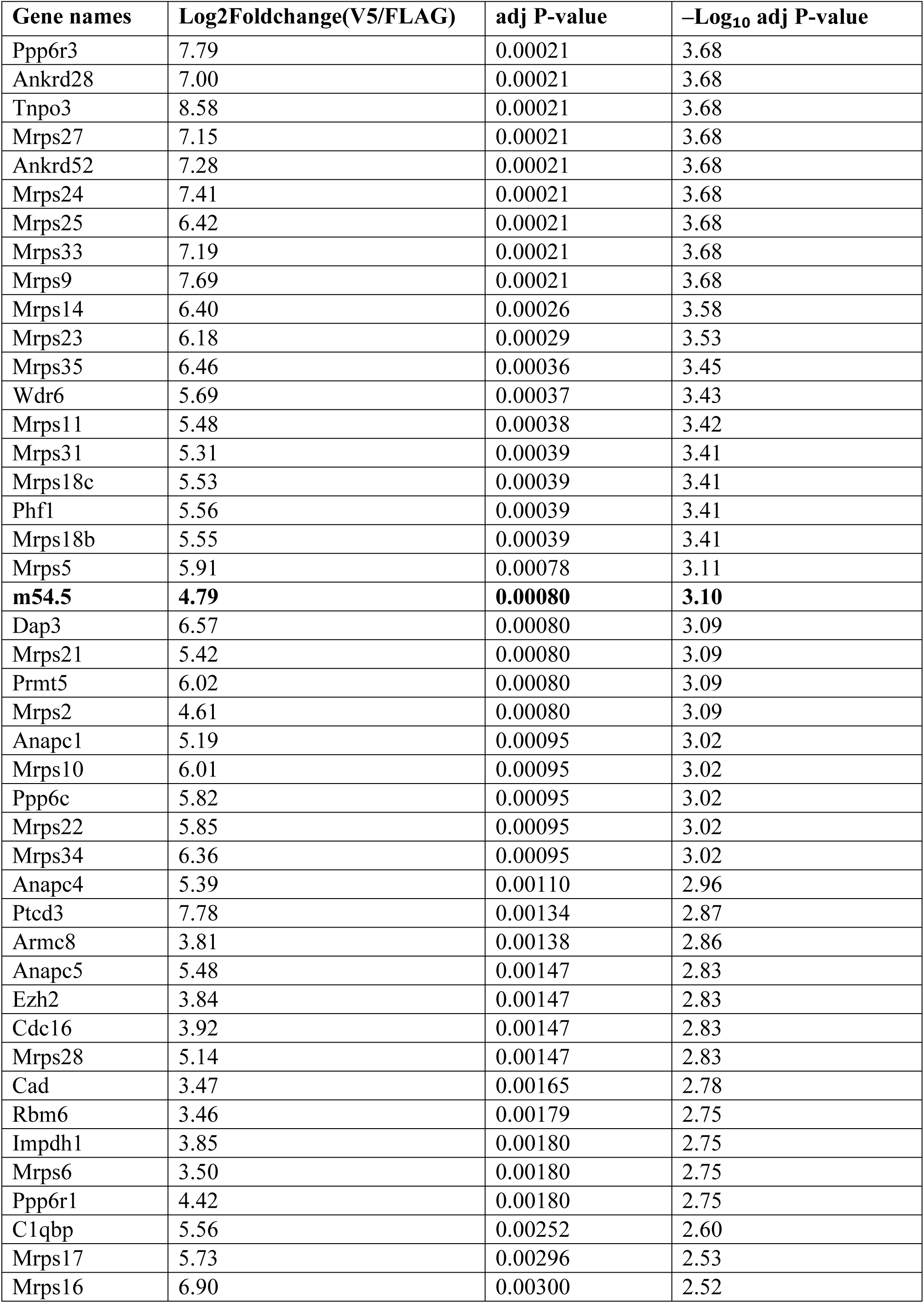
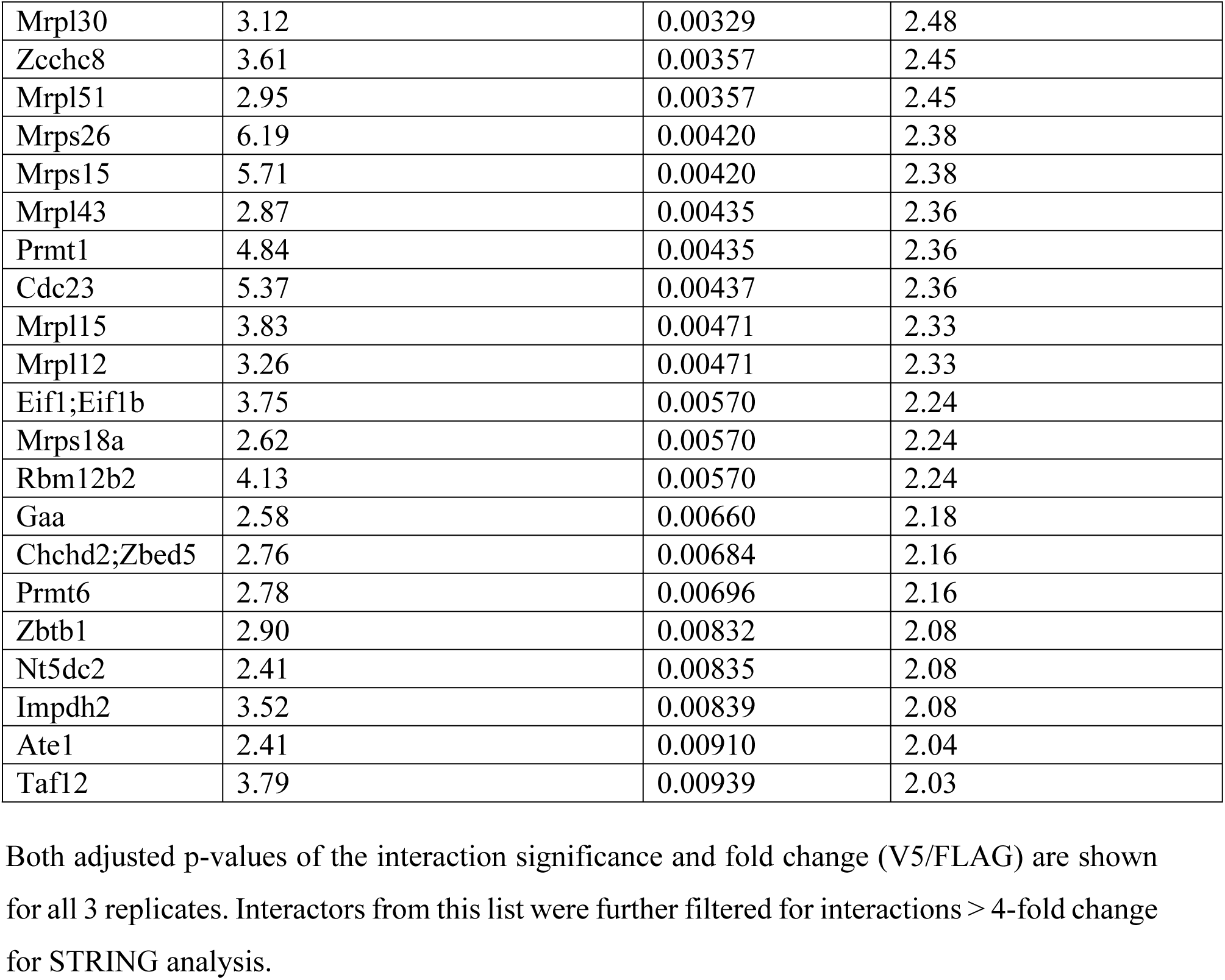
Detailed table of the top interactors of m54.5.

Both adjusted p-values of the interaction significance and fold change (V5/FLAG) are shown for all 3 replicates. Interactors from this list were further filtered for interactions > 4-fold change for STRING analysis.

### m54.5p expression results in cell cycle arrest at the G1/S phase

The APC/C E3 ubiquitin ligase complex is one of the master regulators of the cell cycle. It is modulated by several viruses, including HCMV, which interferes with G1-S phase transition by disrupting the complex [12,19,35]. The PP6 complex has been implicated in regulating mitotic spindle formation and transition into S phase [36], among other functions. We thus asked whether m54.5p indeed regulated the cell cycle. We first analyzed the role of m54.5p in isolation, as MCMV encodes various other cell cycle regulators. We hence generated polyclonal cell lines expressing a doxycycline-inducible C-terminally V5-tagged m54.5p to improve m54.5p expression and prevent host adaptation to the viral protein under constitutive expression. Following cell line validation **(Fig 4A)**, we performed nocodazole mitotic arrest assays by treating doxycycline-pre-treated NIH-3T3 and m54.5p-V5-3T3 (inducible) with nocodazole for 16 hours to cause G2/M phase arrest consistent with tetraploid 4n DNA content. Cells were fixed and stained for DNA content using propidium iodide. Interestingly, while a majority of NIH-3T3 cells treated with nocodazole arrested at G2/M phase (4n), m54.5p-V5-3T3 (inducible) cells showed a significantly greater number of cells arrested at G1 phase (2n) **(Fig 4B, C)**. To investigate the potential impact of G1 cell cycle arrest on S-phase progression, we conducted a serum starvation S-phase assay **(Fig 4D)**. Briefly, NIH-3T3 and m54.5p-V5-3T3 (inducible) cells, pre-treated with doxycycline, were synchronized at the G1 phase by 48 hours of serum starvation. Following synchronization, the medium was replaced with serum-containing medium to induce cell cycle re-entry. S-phase progression was subsequently monitored using EdU labeling coupled with click chemistry. Strikingly, doxycycline-treated NIH-3T3 cells exhibited a significantly higher proportion of S-phase cells compared to similarly treated m54.5p-V5-3T3 (inducible) cells **(Fig 4E, F)**.

**Fig 4:**
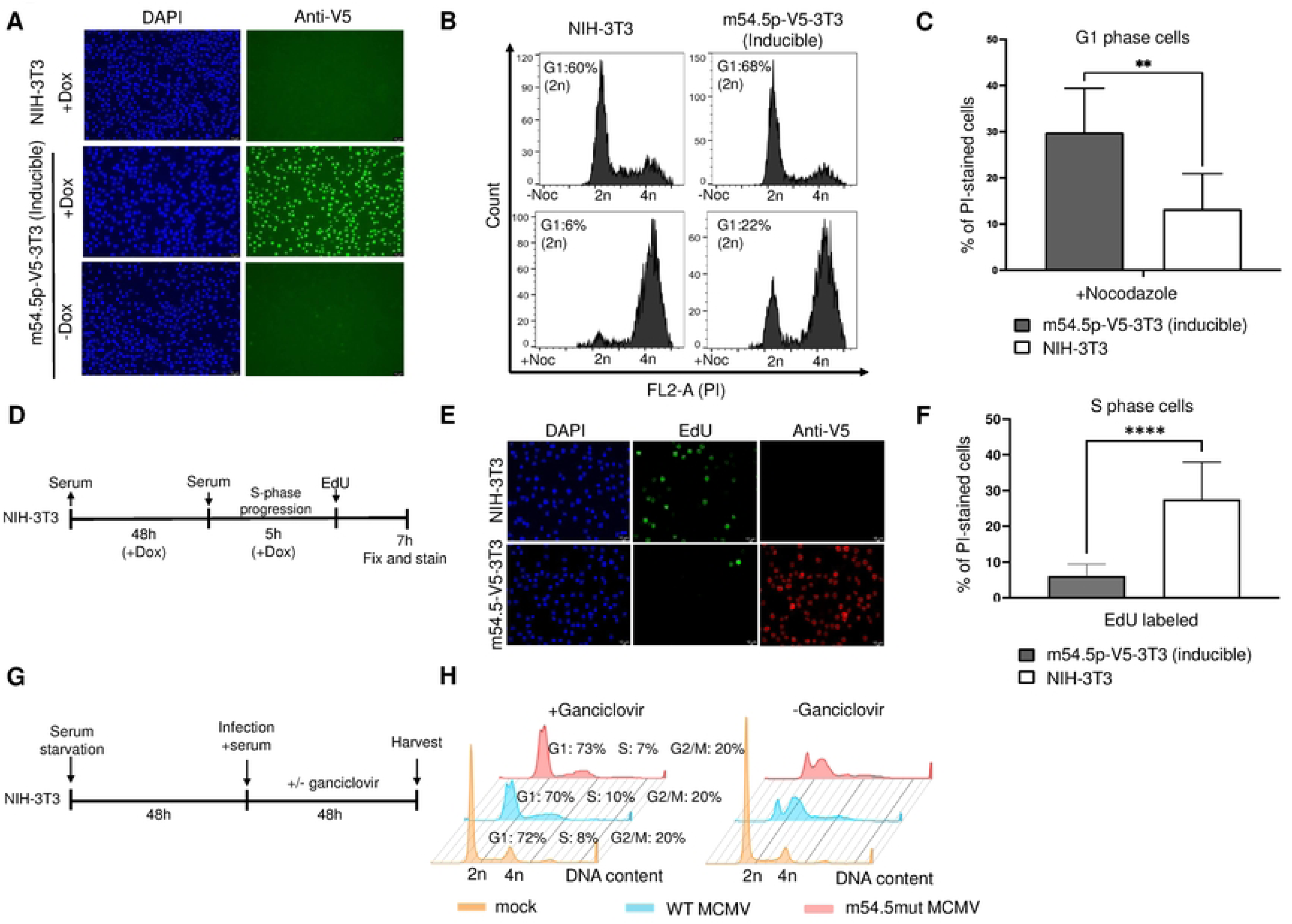
m54.5p expression results in cell cycle arrest at the G1/S phase but exhibits redundancy. **A.** m54.5p-V5-3T3 (inducible) and NIH-3T3 cells treated with or without 1μg/ml doxycycline (Dox) for 24 h were stained for V5. DAPI was used for nuclear counterstaining. **B.** m54.5p-V5-3T3 (inducible) and NIH-3T3 cells treated with 1μg/ml doxycycline (Dox) for 24 h followed by +/− nocodazole (Noc) treatment for 16 h (0.8 μg/ml) in the presence of doxycycline were harvested, fixed, and stained for DNA content via propidium iodide staining in the presence of RNase A using flow cytometry. The percentage of cells in G1 (2n DNA content) is shown for a representative of three biological replicates. **C.** Bar graph showing the percentage of cells in G1 (2n DNA content) from B. Statistical significance was determined using the two-tailed parametric t-test. **D.** Schematic of the serum starvation S-phase assay. **E.** m54.5-V5-3T3 and NIH-3T3 cells were subject to the serum starvation S-phase assay and stained for V5 and newly synthesized DNA via EdU labelling. DAPI was used for nuclear counterstaining. Data from three biological replicates are shown. **, p<0.01. **F.** Bar graph showing the percentage of S-phase cells from E. Statistical significance was determined using the two-tailed parametric t-test. **G.** Schematic of the serum starvation infection assay. **H.** Synchronized NIH-3T3 cells were infected with WT MCMV or m54.5mut MCMV (MOI= 3) in 10% NCS at an MOI of 3. After 1 hour, cells were washed and cultured in serum-free medium with or without 50 µM ganciclovir. After 48 h, cells were fixed, permeabilized, and stained with anti-IE1 antibody (CAPRI HR-MCMV-12) and propidium iodide for flow cytometry analysis of viral m54.5 expression and DNA content. Histogram peaks represent DNA content in IE1-positive gated cells, with 2n DNA content indicating G1 phase and 4n DNA content indicating G2/M phase. The percentage of cells in each cell cycle phase is indicated numerically.

To determine whether similar effects could be observed in virus-infected cells, NIH-3T3 cells were synchronized at the G1 phase through 48 hours of serum starvation and subsequently infected with either WT MCMV or *m54.5mut* MCMV at a multiplicity of infection (MOI) of 3 for an additional 48 hours. To assess the potential impact of m54.5 on host cellular DNA synthesis, viral DNA replication was inhibited using Ganciclovir. Following infection, cells were fixed and immunostained for the viral IE1 protein to identify infected cells, while DNA content was analyzed using propidium iodide staining and flow cytometry **(Fig 4G)**. Notably, no significant difference was observed in the proportion of G1-phase cells infected with WT MCMV compared to *m54.5mut* MCMV **(Fig 4H)**. Based on these findings, we conclude that MCMV m54.5p functions as a cell cycle regulator that impedes S-phase progression by inducing G1-phase arrest in isolation yet exhibits redundancy in its modulation of the cell cycle in MCMV-infected fibroblasts and is thus not required for lytic infection of fibroblasts in vitro.

## Discussion

In this study, we characterized the non-canonical MCMV protein, m54.5p, encoded by a previously unannotated ORF overlapping with the viral polymerase (M54). Our findings demonstrate that m54.5p is an early nuclear protein that interacts with key cell cycle regulators, including the anaphase-promoting complex/cyclosome (APC/C), and induces G1/S phase arrest in murine fibroblasts. Despite its dispensability for viral replication *in vitro*, m54.5p may play a nuanced role in modulating the host cell cycle to facilitate viral genome replication, potentially through functional interactions with the APC/C complex.

The fact that m54.5p has co-evolved within the highly conserved viral DNA polymerase gene (M54) is remarkable. It highlights the high plasticity and dynamic nature of cytomegalovirus genomes and their ability to accommodate novel gene products. Interestingly, both transcriptomics analysis and ribosome profiling data indicated that m54.5p is more abundantly expressed than the conserved viral DNA polymerase M54 expressed within the same genomic locus. Our findings raise important evolutionary questions regarding the conservation of m54.5p and its functional relationship with the viral DNA polymerase. Although m54.5p lacks sequence homology with other cytomegalovirus proteins, its overlapping genomic location with M54 suggests potential regulatory roles in viral DNA synthesis, warranting future studies.

Our results demonstrate that m54.5p expression alone is sufficient to induce G1/S phase arrest, as evidenced by nocodazole and serum starvation assays. This is consistent with findings on HCMV pUL21a [24] and aligns with the well-documented strategy of cytomegaloviruses to manipulate the host cell cycle, creating an S-like environment conducive to viral DNA replication while suppressing cellular DNA synthesis. The *m54.5mut* MCMV mutant exhibited no significant replication defect in fibroblasts, suggesting functional redundancy among MCMV’s cell cycle modulators (e.g., with IE3, M97, and M117). This redundancy highlights the complexity of viral-host interactions and the ability of MCMV to employ multiple strategies to concertedly and optimally achieve the same goal. The non-essentiality of m54.5p *in vitro* contrasts with its conservation across MCMV strains, suggesting a selective advantage *in vivo,* potentially in specific cell types or during immune evasion, like M117 [18].

m54.5p’s interaction with the APC/C complex is particularly compelling, as this E3 ubiquitin ligase is a master regulator of cell cycle progression [37–39]. The APC/C controls the degradation of key cyclins and inhibitors (e.g., cyclin A, cyclin B, and securin) to ensure proper mitotic exit and G1 phase maintenance[40–42]. HCMV disrupts APC/C activity via pUL21a, which exhibits E3 ubiquitin ligase activity and promotes cyclin A degradation and prevents mitotic exit [24]. While MCMV lacks a pUL21a homolog, our findings suggest that m54.5p evolved to functionally converge on the APC/C pathway to enforce G1/S arrest. This underscores the evolutionary adaptation of cytomegaloviruses to exploit host cell cycle machinery, despite differences in the specific viral proteins involved. The interaction with ANAPC1 (APC1), a core scaffold subunit of the APC/C, raises the possibility that m54.5p either stabilizes or inhibits the complex to prolong G1 phase—a strategy that would free nucleotide pools for viral genome replication. Given the strict species specificity of cytomegaloviruses, the independent evolution of m54.5p and pUL21a in MCMV and HCMV, respectively, highlights the importance of cell cycle manipulation for cytomegaloviruses.

Beyond the APC/C, m54.5p also associates with the PP6 phosphatase complex, which has roles in mitotic spindle formation and DNA damage responses [36,43]. PP6 has been implicated in dephosphorylating critical cell cycle regulators, including Aurora kinases and the retinoblastoma (Rb) protein [44–46]. Given that MCMV already targets Rb-E2F pathways via M117 [18], m54.5p’s interaction with PP6 may reinforce cell cycle arrest by modulating phosphorylation-dependent signaling. Alternatively, PP6 could influence viral replication indirectly by regulating stress responses or innate immune signaling. This dual interaction with APC/C and PP6 highlights the multifaceted approach employed by MCMV to manipulate the host cell cycle, ensuring that cellular resources are diverted toward viral replication.

The species-specific nature of m54.5p, with no apparent homologs identified in other CMVs or cellular proteins, suggests that it uniquely evolved in MCMV. While our study provides valuable insights into the role of m54.5p in cell cycle regulation, several limitations should be acknowledged. First, the *in vitro* nature of our experiments may not fully capture the complexity of MCMV infection *in vivo*. Given that the cellular environment and immune pressures differ between cultured fibroblasts and *in vivo* infection, m54.5p may play a more critical role under more natural conditions. In the absence of comprehensive information on MCMV regulators of the cell cycle, we thus decided not to study the function of m54.5p *in vivo*. Second, the redundancy observed in cell cycle modulation by MCMV suggests that additional viral proteins may compensate for the loss of m54.5p, complicating the interpretation of its role, or the cell cycle functions may become more evident in other cell lines. Indeed, M117 exhibits differences in its importance *in vitro* and *in vivo* [18]. Further studies are needed to identify these compensatory mechanisms and their relative contributions to cell cycle regulation.

In conclusion, our findings reveal a surprising plasticity of cytomegalovirus genomes for evolving new viral regulators even in the most deeply conserved viral gene loci. They underscore the importance of viral cell cycle manipulation and highlight surprisingly similar needs of evolutionary distinct cytomegaloviruses. Accordingly, despite millions of years of independent coevolution with their specific hosts, human and mouse CMV independently evolved highly similar means to target the cell cycle during productive infection.

## Materials and Methods

### Cell culture, viruses, and infection

NIH-3T3 (ATCC CRL – 1658) Swiss murine embryonic fibroblasts were grown in DMEM (Dulbecco’s Modified Eagles’ Medium) supplemented with 10% NCS (New-born calf serum) and 100 IU/ml penicillin and 100 μg/ml streptomycin. NIH-3T3 cells engineered to express m54.5 (constitutive/inducible) were generated by transducing the respective lentivirus supernatants using centrifugation at 800G/30 minutes with 1 μg/ml polybrene. Transduced NIH-3T3 cells were grown for three days before selection with 5 μg/ml puromycin to generate polyclonal cell lines. 293T Human embryonic kidney (HEK) epithelial cells were grown in DMEM (Dulbecco’s Modified Eagles’ Medium) supplemented with 10% FCS (Fetal calf serum) and 100 IU/ml penicillin and 100 μg/ml streptomycin. All cells were grown at 37°C at 5% CO_2_ under humid conditions. All MCMV viruses were based on the pSM3fr MCMV Smith background. Crude virus stocks were generated in NIH-3T3 cells. Viruses were titrated by plaque assay on NIH-3T3 cells [47,48]. Cells were infected with MCMV at an MOI of 1 for 1 hour at 37 °C at 5% CO2 with intermittent shaking, followed by media exchange, marking the 0-hour post-infection time-point.

### Virus mutagenesis

The *m54.5mut* MCMV mutant was generated on the pSM3fr MCMV Smith backbone cloned as a bacterial artificial chromosome (BAC) in *E. coli* GS1783 using *en passant* markerless mutagenesis. Briefly, a Kan^r^ marker containing PCR product flanked by homologies to the desired regions in the m54.5 locus harboring a start codon mutation (ATG> TTG) was generated using the CloneAmpHiFi PCR premix (Takara Bio. 639298) using primers 5’- TTGGTGCACTGCTTCATCGCCTCCCGGACCAGCCCAGCTTGCGTTGGGTTAGCCAACGTACAGCAGCTCCCAGCTTGCGTTGGGTTAGCCAACGTACCAGCAGCTCGGCCAGCAGAGAGCGATAGGGATAACAGGGTAATCGATTT-3’ and 5’- TTTGTGCGGGAGAACGTACATCGCTCTCTGCTGGCCGAGCTGCTGGTACGTTGGC ACGTTGGCACGTTGGCTAACCCAACGCAAGCTGGTCCGGGAGGTATATCTGGCCCGTACATCGATCT-3’. The product was transformed into *E. coli* GS1783 containing the MCMV BAC. Following Kan^r^ cassette removal and homologous recombination, selected BAC clones were verified via restriction digestion, and the desired mutation was analyzed via Sanger sequencing. MCMV BAC DNA was purified using the NucleoBond BAC 100 kit (Macherey-Nagel #740579) followed by transfection into NIH-3T3 or m54.5p-V5-3T3 cells using the TransIT-X2 system (Mirus). Upon 100% CPE, virus was passaged twice before producing virus stocks.

### Production of m54.5 antibody

A 6XHis-tagged m54.5 ORF was cloned into the IPTG-inducible pET22b (+) vector (Novagen) via In-fusion cloning (Takara Bio In-fusion HD Cloning Plus kit). The PCR product was generated by using WT MCMV BAC as a template using primers 5’- GTGGTGCTCGAGTGCGGCCGCCATGGGATACCGTTCTTCTTGA-3’ and 5’- CCAGCCGGCGATGGCCATGGATATGGCTAACCCAACGCAAG-3’. The cloned vector was transformed into *E.coli* BL21. m54.5p recombinant protein expression and isolation was performed as per the Qiagen QIAexpressionist. The isolated protein was purified via affinity chromatography on Ni-sepharose columns. Purified protein was then used to immunize BALB/c mice to isolate mouse monoclonal antibodies as described previously. Clones were tested using ELISA on purified m54.5p and the obtained crude antibodies were finally tested via immunoblotting to confirm m54.5p in WT MCMV infection and in m54.5p-V5-3T3 cell lines as a positive control.

### Generation of m54.5p-expressing cell lines

m54.5p-expressing constitutive cell lines tagged with either a 1X V5 or FLAG epitope at the C-terminus were generated by In-fusion cloning (Takara Bio In-fusion HD Cloning Plus kit) of the suitable PCR products into the pMSCV-puro^R^ backbone (digested with EcoRI and XhoI). PCR products were amplified using primers 5’-CCGGAATTAGATCTCTCGAGGCCACCATGGCTAACCC-3’ and 5’- TCCCCTACCCGGTAGAATTCTCATGTACTGTCCAGTCCCAG-3’ (V5 tag) or 5’- TCCCCTACCCGGTAGAATTCTCACTTATCATCGTCATCCTTGTAATC-3’ (Flag tag). The m54.5p-expressing doxycycline inducible sytem was generated by cloning a PCR product into the TET-ON pLT3GG system (lacking the mirE sequence) between BamHI and EcoRI restriction sites. The PCR product was generated by primers 5’- GTCGAGCTTGCGTTGGATCCATGGCTAACCCAACGCAAG-3’ and 5’- CAAGATAATTGCTCGAATTCTCATGTACTGTCCAGTCCCAGG-3’. All PCR products were generated using CloneAmpHiFi PCR premix (Takara Bio. 639298). The obtained plasmids were transformed into Takara Stellar competent cells. Correct clones were analyzed via restriction digestion and Sanger Sequencing. DNA was purified using the ZymoPURE II Plasmid Midiprep kit. Lentiviruses were generated by transfecting HEK293T cells (6-well plate) with one of the above plasmids along with pVSVg and psPax2 (obtained from Addgene) using TransIT-X2 system (Mirus). Lentivirus supernatant (1-2 ml) was harvested 48 hours’ post-transfection to transduce NIH-3T3 cells.

### Immunoblotting

Cells were lysed with 2X Laemmli sample buffer (Cold Spring Harbor protocols). Lysed samples were heated at 95°C/10 minutes. Tris-Glycine SDS-PAGE (8-12%) and wet transfer (Tris-Glycine-20% Methanol) on 0.2 μm Nitrocellulose membrane (Amersham Protran) were performed using the Mini Gel Tank (Life technologies). Membranes were subsequently subject to blocking in 5% (v/v) skimmed milk in 1X PBST (Phosphate buffered saline– 0.1% Tween 20) at room temperature for one hour. Membranes were probed with primary antibodies dissolved in 3% BSA-PBST (0.1% Tween-20), overnight at 4°C at the following concentrations: mouse anti-m54.5p (1:2), rabbit Anti-V5 tag (D3H8Q) mAb (1:1000, Cell Signalling Technology 13202S), rat Anti-DYKDDDDK (FLAG) epitope (L5) antibody (1:1000, Novus Biologicals NBP106712), mouse anti-m123/IE1 (MCMV) antibody – clone IE1.01 (1:1000, CAPRI HR-MCMV-12), mouse anti-M55/gB (MCMV) antibody – clone M55.02 (1:1000, CAPRI HR-MCMV-14), mouse anti-M112-113/E1 (MCMV) antibody – clone CROMA103 (1:1000, CAPRI HR-MCMV-07), mouse Beta Actin antibody (C4) (1:1000, SCBT sc-47778), rabbit α-Tubulin Antibody (1:1000; #2144 Cell Signaling Technology), APC1 Polyclonal antibody (1:1000, 21748-1-AP Proteintech), PPP6C Polyclonal antibody (1:1000, 15852-1-AP Proteintech), rabbit SAPS3 antibody (1:1000, NBP2-34049, Novus Biologicals). Membranes were then washed for 10 minutes thrice in 1X PBST followed by incubation with secondary antibodies dissolved in 3% BSA-PBST (0.1% Tween-20), for 1 hour at RT at the following concentrations: goat Anti-rabbit IgG (Whole molecule – HRP conjugated) (1:10,000, Sigma Aldrich A0545), rabbit Anti-mouse IgG (Whole molecule – HRP conjugated) (1:10,000 Sigma Aldrich A9044), rabbit Anti-rat IgG (Whole molecule – HRP conjugated) (1:10,000 Sigma Aldrich A5795), IRDye 800CW Goat Anti-mouse IgG (1:1000, LI-COR Biosciences 92632210), IRDye 680RD Goat α-rabbit IgG (1:1000, LI-COR Biosciences 92668071). Followed by washing, proteins were analyzed by visualizing the membranes on LI-COR Odyssey FC Imaging System. Proteins bound by HRP-conjugated antibodies were developed using the Thermo Fischer SuperSignal West Pico PLUS substrates as per the company’s instructions.

### Co-immunoprecipitation

V5 Co-immunoprecipitation (Co-IP) samples were prepared from the m54.5p-expressing constitutive cell lines (V5/FLAG-tagged). For each sample, ten million cells were washed and collected by scraping cells in 1ml 1X ice-cold PBS. After centrifugation (1000G/10 minutes), cells were lysed in 1 ml IP lysis buffer (50 mM Tris-HCL (pH:7.4), 300 mM NaCl, 1% (w/v) Triton X-100, 1 mM EDTA, 1 mM PMSF, cOmplete protease inhibitor cocktail (1 tablet/10 mL), Benzonase (50 units/mL, ChemCruz)) for 1 hour at 4°C with constant rotation. Lysed cells were then centrifuged at 18,000 G/15 minutes. The pellet was discarded and 50 μl of supernatant was collected as input. 950 μl was incubated with 25 μl of V5-Trap Magnetic Agarose beads (Chromotek) overnight at 4°C with constant rotation. Beads were magnetically separated using the DynaMag2 (Invitrogen) set up and supernatant was collected as the flow through fraction. Both input and flow through fractions were lysed in 4X Laemmli buffer (2X final concentration). Beads were washed 5 times in IP wash buffer (50 mM Tris-HCL (pH:7.4), 150 mM NaCl, 1% (w/v), cOmplete protease inhibitor cocktail (1 tablet/10 mL)) and finally boiled in 100-200 μl 2X Laemmli buffer at 95°C for 10 minutes to extract proteins (output). Laemmli buffer without DTT was used to make samples for mass spectrometry. Co-IP samples were then subject to Immunoblotting for further analysis.

### Mass spectrometry

Samples lysed in 2X Laemmli buffer were incubated with 4x volume acetone at 20°C/overnight to precipitate proteins. Pellets were washed in acetone at −20°C. Precipitated proteins were dissolved in NuPAGE LDS sample buffer (Life Technologies), reduced with 50 mM DTT at 70 °C for 10 minutes and alkylated with 120 mM iodoacetamide at room temperature for 20 minutes. Separation was performed on NuPAGE Novex 4-12 % Bis-Tris gels (Life Technologies) with MOPS buffer according to manufacturer’s instructions. Gels were washed three times for 5 min with water and stained for 60 min with Simply Blu Safe Stain (Life Technologies). After washing with water for 1 h, each gel lane was cut into 15 slices. The excised gel bands were destained with 30 % acetonitrile in 0.1 M NH_4_HCO_3_ (pH 8), shrunk with 100 % acetonitrile, and dried in a vacuum concentrator (Concentrator 5301, Eppendorf, Germany). Digests were performed with 0.1 µg trypsin per gel band overnight at 37 °C in 0.1 M NH_4_HCO_3_ (pH 8). After removing the supernatant, peptides were extracted from the gel slices with 5 % formic acid, and extracted peptides were pooled with the supernatant. NanoLC-MS/MS analyses were then performed on an Orbitrap Fusion (Thermo Scientific) equipped with a PicoView Ion Source (New Objective) and coupled to an EASY-nLC 1000 (Thermo Scientific). Peptides were loaded on a trapping column (2 cm x 150 µm ID, PepSep) and separated on capillary columns (30 cm x 150 µm ID, PepSep) both packed with 1.9 µm C18 ReproSil and separated with a 30-minute linear gradient from 3% to 30% acetonitrile and 0.1% formic acid and a flow rate of 500 nl/min. Both MS and MS/MS scans were acquired in the Orbitrap analyzer with a resolution of 60,000 for MS scans and 30,000 for MS/MS scans. HCD fragmentation with 35 % normalized collision energy was applied. A Top Speed data-dependent MS/MS method with a fixed cycle time of 3 s was used. Dynamic exclusion was applied with a repeat count of 1 and an exclusion duration of 30 s; singly charged precursors were excluded from selection. The minimum signal threshold for precursor selection was set to 50,000. Predictive AGC was used with AGC a target value of 4×10^5^ for MS scans and 5×10^4^ for MS/MS scans. EASY-IC was used for internal calibration.

### Mitotic arrest assay and flow cytometry

Doxycycline-induced m54.5p-V5-3T3 (inducible) cell lines and NIH-3T3 cells treated with or without Nocodazole (800 ng/ml) for 16 hours were washed in 1X PBS and harvested by trypsinization. Harvested cells were fixed and permeabilized in ice-cold 80% methanol and stored at 4°C for 20 minutes. Fixed cells were washed twice in 1X PBS and incubated with 50 µl RNAse A (100 µg/ml) and 200 µl propidium iodide solution (50 µg/ml) for 30 minutes at 37°C (dark) to stain DNA. Cells were directly strained and used for flow cytometry analysis using BD FACSCalibur. Briefly, NIH-3T3 Cells were gated via FSC vs SSC analysis, followed by segregation of single cells by FL-2 area (A) vs width (W) channels. The FL2-H channel was then utilized to study propidium iodide-stained cells and cell cycle.

### Immunofluorescence

Seeded cells were washed in 1X PBS and fixed in 4% paraformaldehyde for 15 minutes at RT, followed by permeabilization in 0.5% Triton-X 100 for 5 minutes at RT. Cells were washed twice in 1X PBS followed by blocking in 10% FCS-1X PBS. Primary antibodies were diluted in 10% FCS-1X PBS in the following concentrations: Mouse Anti-m54.5p (1:2), Rabbit Anti-V5 tag (D3H8Q) mAb (1:1000, Cell Signalling Technology 13202S), Rat Anti-DYKDDDDK (FLAG) epitope (L5) antibody (1:1000, Novus Biologicals NBP106712). Cells were washed twice in 1X PBS and incubated with secondary antibodies (diluted in 10% FCS-1X PBS) in the following concentrations: Anti-Rabbit IgG (H+L) AlexaFluor 647 (1:1000, Thermo Fisher Scientific A21247), Anti-Rat IgG (H+L) AlexaFluor 488 (1:1000, Thermo Fisher Scientific A11006), Anti-Mouse IgG (H+L) AlexaFluor 568 (1:1000, Abcam ab175473). Cells were washed twice in 1X PBS and incubated with 1X DAPI solution for 5 minutes to stain nuclei before washing and processing. Microcopy was conducted using the Leica DMi8 (Leica Microsystems) device as per the manufacturers’ instructions.

### Edu Labelling and click chemistry

NIH-3T3 cells were labelled with 10 µM EdU for a duration of 1 hour to label DNA-synthesizing cells. Following fixation and Permeabilization, click reaction was conducted using sodium ascorbate (10mM), CuSO_4_ (1mM), amino guanidine (10mM), and thiamine triphosphatase (THPTA (1mM)) along with clickable azide (488 nm – 10 µM) for 60 minutes at RT in the dark. Cells were then washed thrice in 1X PBS and processed for further staining (immunofluorescence) or microscopy.

### Serum starvation-infection assay

NIH-3T3 cells were synchronized by serum starvation for 48 hours and subsequently infected with either WT MCMV or *m54.5mut* MCMV in the presence of 10% newborn calf serum (NCS). After 1 hour of infection, cells were washed, and fresh serum-free medium supplemented with 50 µM ganciclovir was added. To simultaneously assess DNA content and viral protein expression, the cells were fixed and permeabilized in 80% ethanol on ice for a minimum of 30 minutes. Following fixation, the cells were immunostained using an anti-IE1 antibody (CAPRI HR-MCMV-12) and a secondary anti-mouse IgG-Alexa405 antibody. DNA content was evaluated by staining with propidium iodide, and samples were analyzed by flow cytometry.

### Data analysis and statistics

All immunoblotting data were analyzed on ImageStudio Lite (LI-COR). Flow cytometry analysis was performed on CellQuestPro (BD Biosciences) during data acquisition and FlowJo 10 (BD Biosciences) to perform gating as well as histogram analysis. Sanger Sequencing data were analyzed on Snapgene (Dotmatics). Graphs were plotted using GraphPad Prism 9 (Dotmatics) unless otherwise stated. Student’s paired two-tailed parametric t-test was used to determine statistical significance and p-values. For MS data, raw MS data files were analyzed with MaxQuant version 1.6.2.2[49]. Database search was performed with Andromeda, which is integrated in the utilized version of MaxQuant. The search was performed against the UniProt Mouse Reference Proteome database (March 2022, UP000000589, 21985 entries) and the UniProt Mouse cytomegalovirus Reference Proteome Database (March 2022, UP000122533, 164 entries). The sequence of the m54.5p was added to this database. Additionally, a database containing common contaminants was used. The search was performed with tryptic cleavage specificity with three allowed mis-cleavages. Protein identification was under the control of the false discovery rate (FDR; <1% FDR on protein and peptide spectrum match (PSM) level).

In addition to MaxQuant default settings, the search was performed against the following variable modifications: Protein N-terminal acetylation, Gln to pyro-Glu formation (N-term. Gln) and oxidation (Met). Carbamidomethyl (Cys) was set as a fixed modification. Further data analysis was performed using R scripts developed in-house. LFQ intensities were used for protein quantitation. Proteins with less than two razor/unique peptides were removed. Missing LFQ intensities were imputed with values close to the baseline. Data imputation was performed with values from a standard normal distribution with a mean of the 5% quantile of the combined log10-transformed LFQ intensities and a standard deviation of 0.1. For the identification of significantly enriched proteins, median log2 transformed protein ratios were calculated from the three replicate experiments, and boxplot outliers were identified in intensity bins of at least 300 proteins. Log2 transformed protein ratios of sample versus control with values outside a 1.5x (significance 1) or 3x (significance 2) interquartile range (IQR), respectively, were considered as significantly enriched in the individual replicates. In addition, the R package limma was used to calculate Benjamini-Hochberg adjusted p-values and FDR[50,51].

## Data Availability Statement

All mass spectrometry proteomics data have been deposited to the ProteomeXchange Consortium via the PRIDE partner repository with the dataset identifier **PXD066167**.

## Acknowledgments

L.D. and M.L. designed the experiments. Y.Z. and M.L. performed experiments, analyzed data, and wrote the paper. V.J.L. developed and provided critical reagents. S.L. from A.S.’s group conducted mass spectrometry analysis. All authors reviewed and approved the final manuscript.

## Supporting information

**S1 File. Multiple Sequence Alignment of m54.5 ORF from Diverse Mouse CMV Strains.** Multiple sequence alignment of the M54.5 open reading frame (ORF) from various mouse cytomegalovirus (CMV) strains, generated using Clustal Omega and visualized with Jalview. The start codon (ATG) is marked in green, stop codon (TGA) is shown in black, and all other nucleotides are color-coded according to adenine (A), thymine (T), guanine (G), and cytosine (C) classification. This alignment illustrates 100% sequence conservation of the m54.5 ORF among the available six MCMV strains.

**S2 File. Potential ORFs in Polymerase Genes of MCMV, HCMV, RCMV and GPCMV.** For each viral polymerase gene sequence (MCMV M54, HCMV UL54, RCMV E54, and GPCMV GP54), the DNA was translated in all three forward reading frames. The canonical polymerase frame is displayed at the top, with the two alternative reading frames shown below, and a nucleotide length ruler is presented at the bottom of each panel. Potential open reading frames (ORFs), defined as regions extending from a start codon (AUG) to the next in-frame stop codon, are highlighted with a light-yellow background, and the number of amino acids (aa) encoded by each ORF is indicated. In the M54 frame, putative ORFs of 227, 119, and 73 aa were identified. Ribo-seq only confirmed the 227 aa m54.5 ORF; in UL54, putative ORFs of 98, 80, and 37 aa were identified; in the alternative frames of E54, putative ORFs of 114, 73, 55, and 39 aa were identified; and in GP54, putative ORFs of 54, 52, 43, and 42 aa were identified. Of note, no ORFs of relevant length were identified in the C-terminal parts of the M54 homologs, where the m54.5 ORF is located.

**S3 File. Comparative conservation of m54.5 protein sequences identified by tBLASTn.** The protein sequence of MCMV m54.5 was used as a query in a tBLASTn search against the NCBI nucleotide database. The top 10 significant hits from different cytomegalovirus (CMV) organisms were retrieved as nucleotide sequences, translated into amino acid sequences using EMBOSS Transeq, and aligned with Clustal Omega. The resulting multiple sequence alignment was visualized in Jalview with the “Clustal” color scheme, highlighting conserved residues. Stop codons are indicated by white asterisks (*) on a black background. Sequence conservation across all aligned proteins is indicated using the conservation track in Jalview.

